# Is synuclein aggregation a derived or ancestral trait? Ancestral sequence reconstruction uncovers stepwise evolution of synuclein aggregation

**DOI:** 10.1101/2025.09.27.678990

**Authors:** Andrew Sam, Seungwoo Lee, Jordan Elliott, Emma Larson, Abdullah Naiyer, Jonathan K. Williams, Jean Baum

**Affiliations:** Department of Chemistry & Chemical Biology, Rutgers University, 123 Bevier Road, Piscataway, NJ 08854, USA; Department of Heath Informatics, Rutgers, the State University of New Jersey, Newark, NJ, 07107-1709; Drug Product Development, Bristol Myers Squibb, 1 Squibb Drive, New Brunswick, NJ 08901, USA

## Abstract

Protein aggregation drives many neurodegenerative diseases, including Parkinson’s disease, where misfolded α-synuclein (αSyn) forms fibrillar assemblies that accumulate as Lewy bodies. Although αSyn aggregation has been extensively characterized, its evolutionary origins and sequence determinants remain unresolved. Here, we use ancestral sequence reconstruction (ASR) to trace the emergence of fibril-forming ability in the synuclein family. We inferred synuclein phylogeny and experimentally resurrected common ancestors, including ROOT synuclein, the last common ancestor of all synucleins, and key intermediates along the αSyn lineage. Strikingly, ROOT synuclein is non-aggregating, demonstrating that fibril formation is an evolved, rather than ancestral property. Aggregation first emerges at the ancestral αβ node, is retained in αSyn, and suppressed in β-synuclein. Biophysical analyses including mass spectrometry and NMR reveal that aggregation aligns with greater complexity and heterogeneity in the monomer conformational ensemble, suggesting that evolutionary sequence changes progressively remodel monomer landscapes to favor fibril formation. Complementing these insights, comparative sequence analysis reveals that the transition from ROOT to α-WT is marked by the stepwise acquisition of residues critical for stabilizing the fibril core. Early mutations stabilized the β-arch core, enabling the onset of fibril formation, followed by substitutions that reinforce protofilament–protofilament interactions. Together, ASR defines an evolutionary framework for synuclein aggregation linking progressive sequence evolution and conformational complexity to the molecular origins of αSyn fibril formation.

## INTRODUCTION

Protein aggregation is a defining feature of many neurodegenerative disorders, including Alzheimer’s disease, Huntington’s disease, and Parkinson’s disease (PD). In these diseases, specific proteins misfold and self-assemble into insoluble fibrillar aggregates that accumulate in neurons, disrupting cellular homeostasis and contributing to progressive neurodegeneration. In PD, a central pathological hallmark is the formation of Lewy bodies—large, intracellular inclusions composed primarily of α-synuclein (αSyn)^1,2^. αSyn is an intrinsically disordered protein abundantly expressed in presynaptic terminals, where it is thought to play a role in synaptic vesicle trafficking and neurotransmitter release^3,4^. Under pathological conditions, αSyn undergoes a conformational transition from a soluble monomer to oligomers and amyloid fibrils that are neurotoxic and capable of propagating between cells in a prion-like manner^5,6^. Despite extensive research, the molecular mechanisms by which sequence variation governs αSyn aggregation remain poorly understood. Understanding these mechanisms is critical for elucidating disease etiology and for developing targeted therapeutic strategies.

Ancestral sequence reconstruction (ASR) offers a novel approach to address this gap by experimentally probing the evolutionary origins of aggregation determinants^7^. ASR uses modern protein sequences to infer the most likely ancestral sequences, enabling the experimental resurrection of proteins that existed along the evolutionary lineage^8^. These ancestral proteins provide unique insights into how molecular properties emerged and diversified over time. Unlike conventional mutagenesis, which perturbs one or a few residues at a time, ASR samples broader sequence space by inferring evolutionary intermediates. ASR has been applied to different systems including, the thermotolerance and viability in harsh condition of ancestral Trx enzymes^9^, the evolution of DNA binding specificity in the steroid hormone receptor family^10^, the evolution of protein thermostability and the thermal adaptation of ancestral catalysts^11,12^. Applied to synucleins, ASR allows us to ask whether the ability to form fibrils is an ancestral property or a derived trait, and to pinpoint the mutational steps that confer fibril-forming ability to αSyn.

The synuclein family, comprising αSyn, β-synuclein (βSyn), and γ-synuclein (γSyn), provides an ideal system for such inquiry. All three proteins share a conserved domain architecture: an N-terminal domain (NTD) that is amphipathic, a central hydrophobic non-amyloid component (NAC) domain, and a highly charged C-terminal domain (CTD). Notably, only αSyn aggregates under physiological conditions, whereas βSyn and γSyn resist fibril formation. βSyn, predominantly found in the brain and spinal cord, contains an 11-residue deletion in the NAC domain that inhibits aggregation at neutral pH but not under acidic conditions^13^. γSyn, primarily expressed in the peripheral nervous system, retains a full-length NAC domain but also requires low pH for aggregate formation^14,15^. Moreover, several heritable mutations in the αSyn NTD modulate aggregation kinetics^16-19^ and are linked to PD progression, underscoring the complexity of sequence-dependent aggregation.

In this study, we applied ASR to investigate the evolutionary origins of synuclein aggregation and the sequence determinants that modulate fibril formation. We reconstructed and experimentally characterized ancestral synuclein proteins, including ROOT synuclein, the last common ancestor of all synucleins, and key intermediates along the phylogeny. Notably, ROOT synuclein did not aggregate under physiological conditions, indicating that aggregation propensity is a derived trait that emerged later in synuclein evolution. Aggregation first appeared at the ancestral αβ node (Anc-αβ), the common ancestor of αSyn and βSyn, and was retained in the αSyn lineage while being suppressed in βSyn. In the αSyn lineage, mass spectrometry and NMR revealed that monomer conformational heterogeneity and increasing collisional cross sections correlate with aggregation propensity, while sequence comparisons identified a stepwise accumulation of fibril-stabilizing residues. Together, these findings suggest that synuclein aggregation is a derived trait that emerged through progressive sequence changes that lead to the diversification of structural ensembles. This evolutionary framework connects molecular variation with the origins of amyloid formation and provides a foundation for understanding how protein evolution shapes neurodegenerative disease.

## RESULTS

### Synuclein phylogeny and ancestral sequence reconstruction identify ROOT, the last common ancestor of all synucleins, and intermediate ancestral synucleins

To perform phylogeny inference and ASR, a dataset of αSyn, βSyn, and γSyn sequences was compiled from the NCBI Proteins Database including 603 unique synuclein sequences. The validity of the dataset was assessed by performing a multiple sequence alignment and calculating the consensus sequence and conservation scores for each position of the alignment (Figure S1). The NTD (residues 1-60) of the sequence alignment was found to be highly conserved, including the well-known imperfect “KTKEGV” repeats which are present throughout the N-terminus of the consensus sequence. Furthermore, the significantly diminished alignment conservation score of positions 73-84 of the NAC (residues 61-95) domain accurately reflects the 11-residue deletion seen in βSyn but not αSyn or γSyn sequences, and the consensus sequence accurately represents the high hydrophobicity of the NAC domain. The conservation score of the CTD (residues 96+) and lack of consensus sequence, with the notable exception of numerous glutamate residues, reflects the highly variable and charged state of the C-terminus. High overall consistency of the dataset with established extant synuclein sequence features implies a representative dataset prime for phylogeny construction and ASR.

To elucidate the evolutionary relationships among synuclein proteins, we constructed a phylogeny using Randomized Accelerated Maximum Likelihood (RAxML)^20^ based on our compiled dataset and multiple sequence alignment. The resulting tree resolved into three distinct clades, each containing extant sequences of αSyn, βSyn, or γSyn (Figure 1, and Figure S2). Notably, the γSyn clade displayed the longest branch lengths, reflecting a higher number of sequence changes and greater sequence diversity among extant γSyn proteins. In contrast, the αSyn and βSyn clades exhibited shorter branches, indicating higher sequence similarity within each group. Moreover, the short branch lengths separating αSyn and βSyn from their last common ancestor highlight the close evolutionary relationship between these two synuclein paralogs.

**Figure 1.**
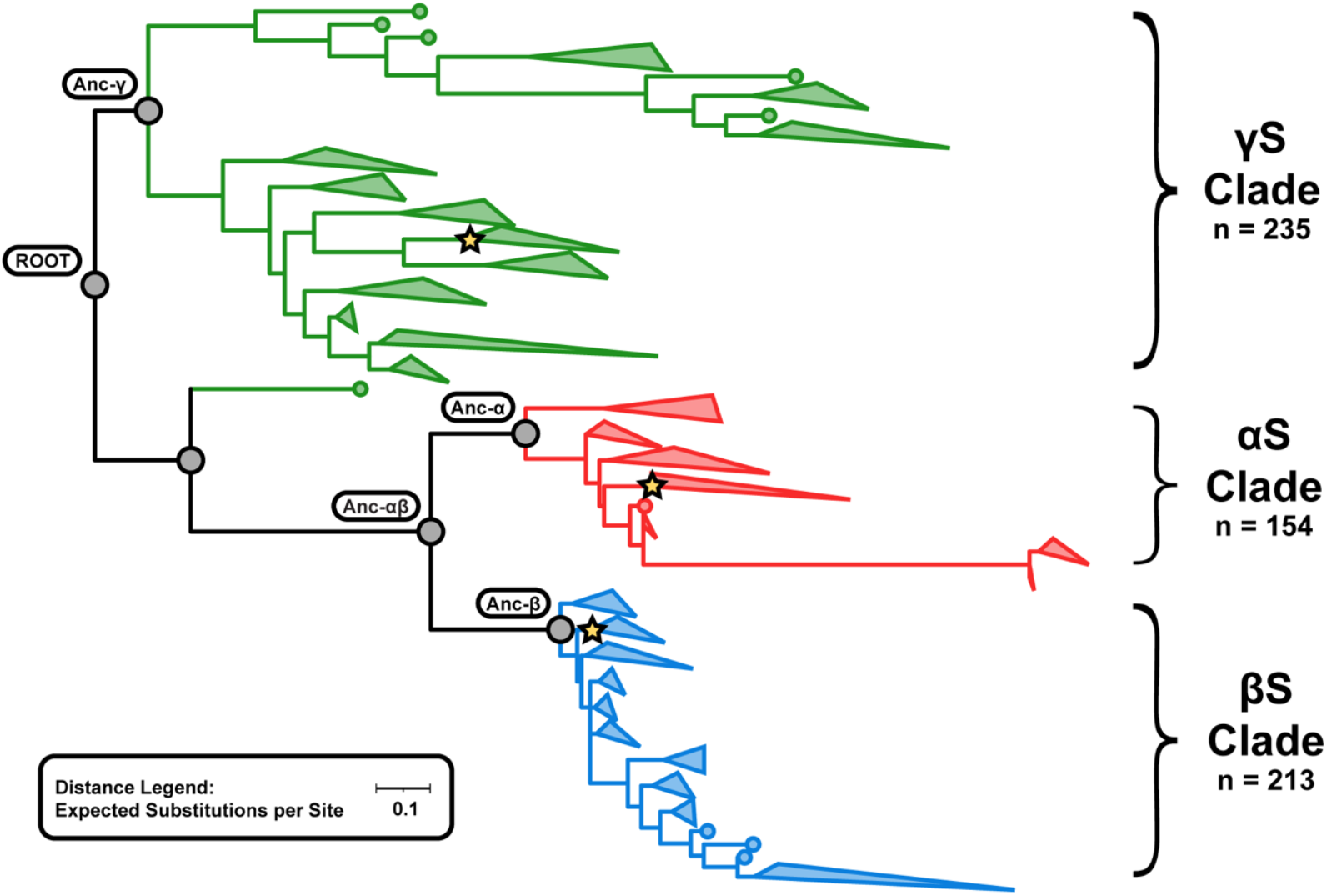
Inferred phylogeny of the synuclein family from ancestral sequence reconstruction. Inferred synuclein phylogeny of a compiled dataset containing 603 distinct protein sequences using ancestral sequence reconstruction. The rooted tree is colored according to synuclein clades, with αSyn (red), βSyn (blue), and γSyn (green). Key ancestral nodes are labeled with a grey circle and extant human αSyn, βSyn, and γSyn are indicated by yellow stars. The total number of extant sequences per clade is indicated. Branch lengths represent genetic distance measured in units of expected substitutions per residue. Branches were collapsed when the average length is less than 0.3 substitutions per site. Scalene triangles illustrate the range of genetic distances between the most closely and most distantly related sequences within each collapsed branch.

Phylogenetic reconstruction suggests that the last common ancestor of all synucleins, ROOT, first diverged into ancestral γ-synuclein (Anc-γ) and ancestral αβγ-synuclein (Anc-αβγ), consistent with an initial gene duplication event^21^. Anc-αβγ then gave rise to Anc-αβ, the most recent common ancestor of extant αSyn and βSyn. A subsequent duplication of Anc-αβ produced Anc-α and Anc-β, establishing the distinct αSyn and βSyn clades. Overall, the features of this phylogeny align with previously reported analyses of synuclein evolution^21,22^, supporting both the robustness of our dataset and the inferred relationships (Figure 1, S2). For experimental characterization, we focused on ROOT, Anc-αβ, Anc-α, and Anc-β, given their direct relevance to extant αSyn and βSyn in neurodegeneration. By contrast, the γSyn clade and Anc-αβγ were not pursued, as extant γSyn is predominantly expressed in peripheral tissues and Anc-αβγ was inferred from only a single extant sequence.

We next compared the reconstructed ancestral sequences to extant human synucleins to identify residues that were retained or lost over evolutionary time. Multiple sequence alignment of ROOT, ancestral nodes, and extant sequences (Figure 2) revealed several key features: high sequence conservation in the NTD and extreme C-terminal regions (the last 10 residues), conservation of the KTKXGX motifs in the N-terminus across all ancestral nodes, and a highly variable CTD. The length of the NAC region in ROOT, Anc-αβ, Anc-α, and α-WT is similar, in contrast to the 11-residue deletion observed in Anc-β and β-WT. A detailed breakdown of residue composition by charge, polarity, and other properties is provided in Figure S3. Specifically, we observed seven mutations between ROOT and α-WT in the N-terminal domain (K5M, F8L, R10K, E31G, M38L, T42S, S53A), 15 mutations in the NAC domain (A63V, N64T, V65N, E68G, S72T, S73G, N75T, T76A, K79Q, E86G, N87S, V88I, V89A, T91A, L94F), and 28 mutations in the CTD (E98D, D99Q, V101G, R102K, P103N, P106E, A107G, E108A, A109P, A110Q, P111E, E117G, Q121I, A122L, G123E, E124D, L125M, G126P, Q127V, A128D, G129P, E130D, G131N, N136M, E137P, G138S, N139E, E142G), in addition to 10 deletions. These results delineate the key sequence changes along the αSyn lineage, setting the stage to connect mutations with the evolutionary emergence of aggregation.

**Figure 2.**
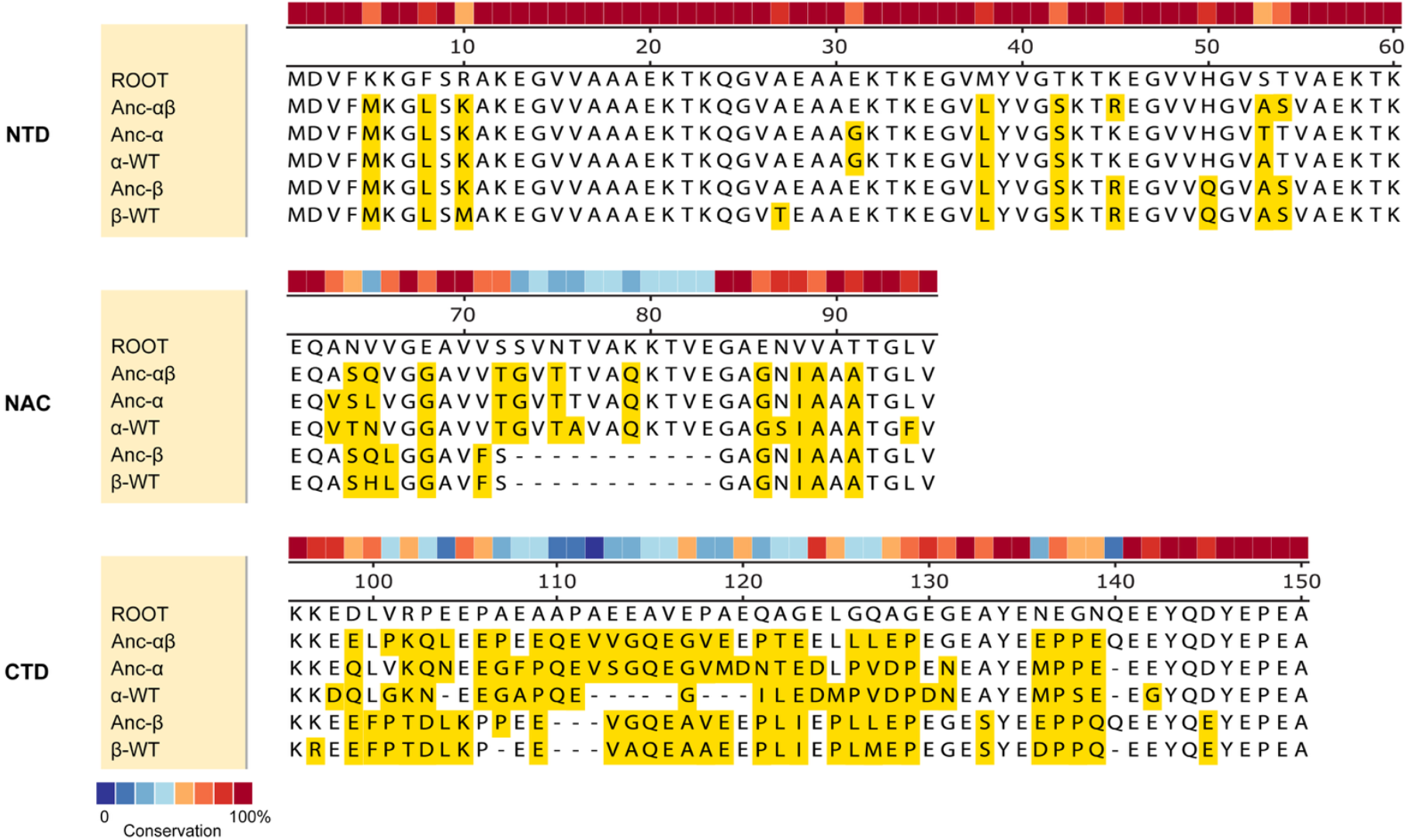
Multiple sequence alignment of extant and ancestral synuclein sequences. Multiple sequence alignment (Clustal Omega) of ROOT, Ancestral-αβ, Ancestral-α, Ancestral-β (inferred from ASR), extant α-WT, and extant β-WT. Sequences are ordered according to evolutionary progression, from top to bottom, and separated by domain (NTD, NAC, CTD). Residues that differ from the ROOT sequence are highlighted in yellow, and dashes indicate gaps from the MSA. Conservation across positions, based on the MSA, is shown by color blocks above the sequences.

### ROOT synuclein resists aggregation; aggregation first emerges at Anc-αβ

To investigate how aggregation evolved across the synuclein family, we measured the fibril forming propensity of ancestral and extant proteins using Thioflavin T (ThT) assays. These experiments reveal two key insights into the evolution of aggregation across synucleins. First, the ROOT sequence does not aggregate under physiological conditions (Figure 3). To further support the intrinsic resistance of ROOT to aggregation, we tested additional conditions known to promote fibril formation, including low pH and higher protein concentrations. ROOT did not aggregate under these conditions (Figure S4) and remained fully soluble (Figure S5), indicating intrinsic resistance to higher order assembly.

**Figure 3.**
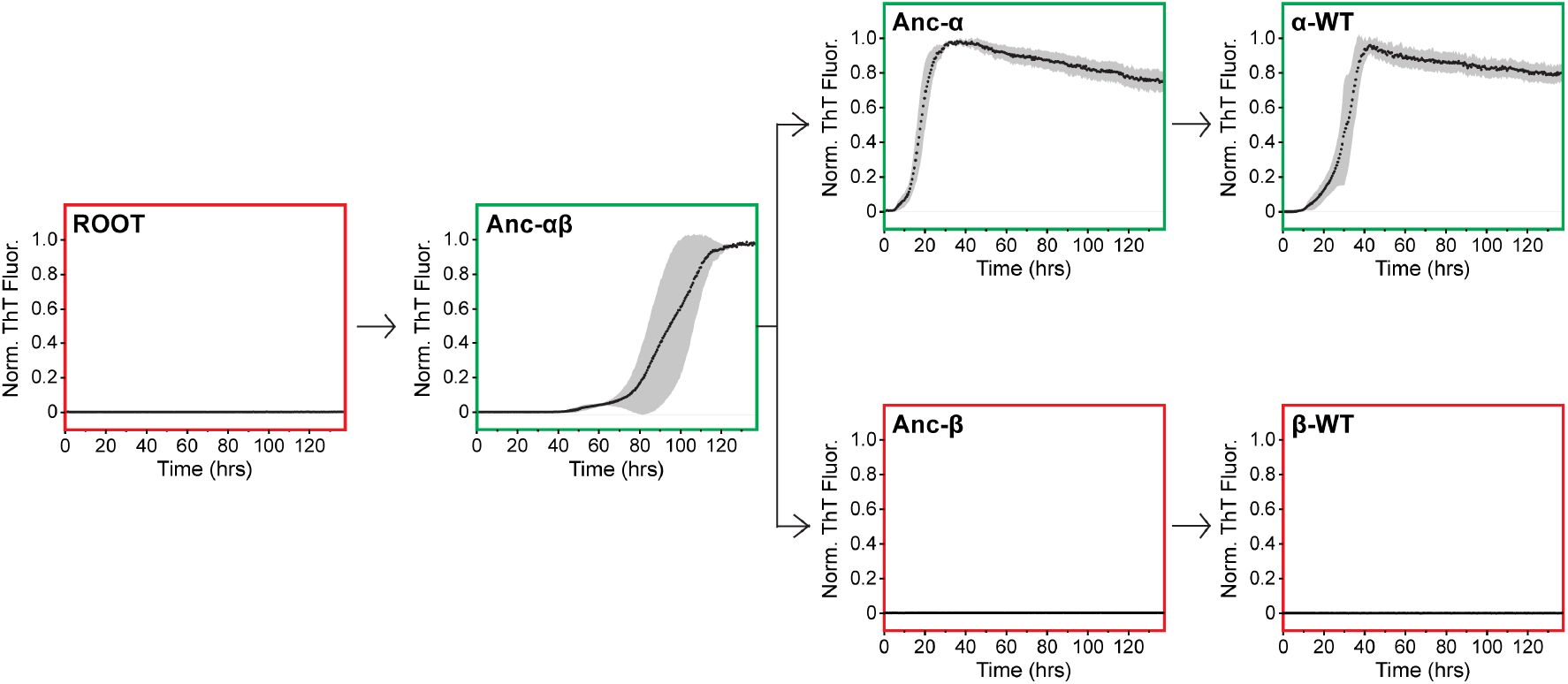
Synuclein aggregation evolved at Anc-αβ, common ancestor of αSyn and βSyn. Comparison of ThT aggregation assays of ROOT, ancestral synucleins, α-WT, and β-WT. Experiments were conducted using 70 μM protein in PBS (pH 7.4), supplemented with a Teflon bead, with shaking at 600RPM at 37°C. Curves represent the mean of six replicates, with shaded grey areas indicating the standard deviation. Green and red boxes indicate species that did and did not aggregate, respectively.

Second, aggregation first emerges at the Anc-αβ node. This node marks the evolutionary divergence of the α-syn lineage toward robust fibril formation, while the β-syn branch, including Anc-β and β-WT, remains largely non-aggregating with nearly all protein remaining soluble (Figure S5). In contrast, the α-syn branch exhibits robust aggregation.

Anc-αβ exhibits a lag time of 72.9 ± 1.0 hrs, approximately threefold longer than α-WT (20.4 ± 0.8 hrs) (Figure 3 and Figure S5). Anc-α aggregates more rapidly than α-WT, with a lag time (11.3 ± 1.1 hrs) nearly half that of α-WT. Residual soluble protein measurements shows that 40.6 ± 0.9 % of Anc-αβ remains monomeric, whereas Anc-α and α-WT display similar levels of residual monomer of approximately 15.3 ± 0.4 % and 11.1 ± 0.2 % respectively (Figure S5). Additionally, atomic force microscopy (AFM) images of the fibril-forming synucleins showed differing morphology, suggesting potentially divergent aggregation mechanisms (Figure S6).

To investigate the sequence determinants underlying the evolution of aggregation from ROOT to Anc-αβ, we generated a series of domain swapped chimeras (Figure 4). We assessed their aggregation propensity (Figure 4 and Figure S5) and found that of the six chimeras, three -RαβR, Rαβαβ, and αβαβR-formed fibrils while the other three-RRαβ, αβRR, and αβRαβ-did not form fibrils. These data suggest that Anc-αβ NAC is essential for aggregation while ROOT NAC inhibits fibril formation. Moreover, kinetic analysis for RαβR, Rαβαβ, and αβαβR revealed that the NTD and CTD modulate aggregation rates, with the Anc-αβ NTD shortening the lag time. Notably, αβαβR exhibited the fastest rate of fibril formation compared to RαβR, Rαβαβ, ancestral and extant proteins, and the lowest residual soluble protein, suggesting a synergistic interaction between the Anc-αβ NAC and the Anc-αβ NTD that enhances aggregation of the αβαβR chimera. Together, these results demonstrate that aggregation resistance is intrinsic to ROOT, that aggregation first arises at Anc-αβ, and that the Anc-αβ NAC domain is necessary for fibril formation.

**Figure 4.**
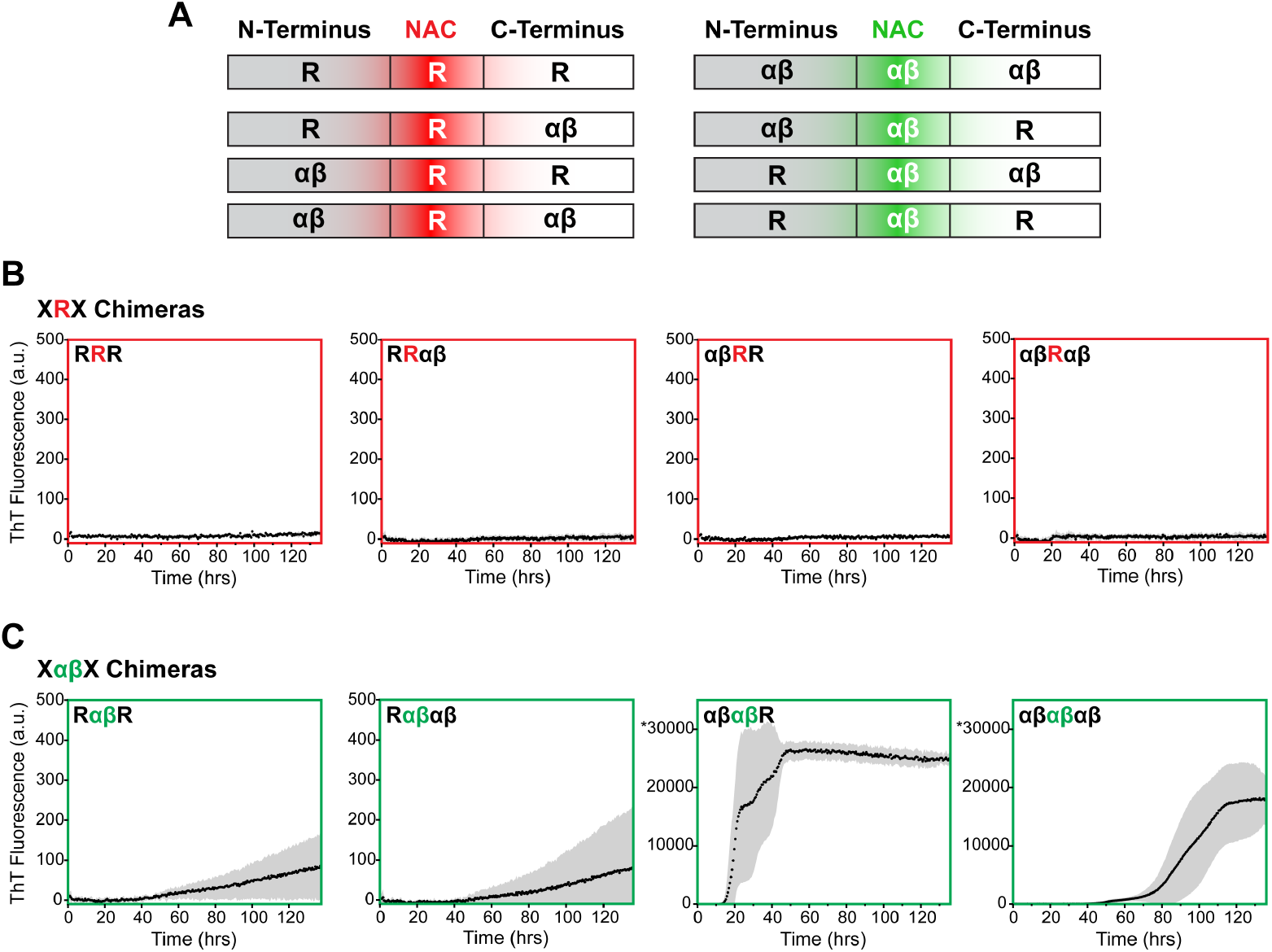
Chimeras containing ROOT NAC resist aggregation, while chimeras containing Anc-αβ NAC promote aggregation. (A) Schematic diagram of domain-swapped chimeras constructed using ROOT and Anc-αβ. ‘R’ represents ROOT sequence and ‘αβ’ represents Anc-αβ sequence. Chimeras containing ‘R’ NAC are highlighted in red and those with ‘αβ’ NAC in green (B) ThT aggregation kinetics of ROOT NAC chimeras at 70 μM in PBS (pH 7.4). (C) ThT aggregation kinetics of Anc-αβ NAC chimeras at 70 μM in PBS (pH 7.4). ThT curves represent averages of at least three independent replicates, with standard deviation indicated in grey. Green and red boxes indicate species that did and did not aggregate, respectively. Asterisks for αβαβR and αβαβαβ indicate altered Y-axis scale due to high fluorescence signal from their strong aggregation propensity.

### Native state ion mobility-mass spectrometry (IMS) highlights ROOT compaction and evolutionary progression in monomer conformational diversity of αSyn lineage

α-WT has been shown to exist in conformational equilibrium among 4 main conformer populations comprised of compact, intermediate compact, intermediate extended, and extended species^23-25^ This conformational diversity, and the degree of molecular compaction, can be probed using native electrospray ionization mass spectrometry (ESI-MS) and ion mobility-mass spectrometry (IMS). When applied to ancestral synucleins, native ESI-MS revealed the characteristic bimodal charge-state distribution, reflecting dominant compact and extended populations (Figure S7) and demonstrating that ancestral synucleins, like the extant synucleins, populate four main populations of compact and extended conformers.

To further analyze the monomer conformations, we employed IMS which can be used to separate conformers and their subpopulations by their extent of compaction. It has been shown that the collision cross section (CCS) of gaseous ions correlate well with those in solution, thus allowing us to use IMS to determine the CCS of each species^26^. The +8 ion is typically used as a fingerprint for analysis of α-WT conformers due to the high conformational heterogeneity and the fact that it sits at an interface between the main extended and compact populations. Comparison of CCS heat maps for +8 ions revealed that ROOT displayed limited conformational heterogeneity and a lower CCS value (2461 Å^2^) relative to Anc-αβ, Anc-α, and α-WT (Figure 5). The α-syn lineage showed progressive increases in CCS and conformational heterogeneity, with α-WT exhibiting four distinct conformations with a mean CCS of 2547 Å^2^. Anc-αβ (2493 Å^2^) displayed an extended distribution of closely spaced conformers, while Anc-α (2683 Å^2^) resolved into three distinct species. In contrast, the β-syn lineage (Anc-β and β-WT) exhibited the lowest CCS values (2367 and 2344 Å^2^, respectively) and restricted heterogeneity, with β-WT dominated by a single conformational state (Figure 5). IMS analysis of chimeras further revealed that aggregating constructs adopted more extended conformations (2557–2656 Å^2^), whereas non-aggregating constructs were more compact (2507–2575 Å^2^) (Figure S8). These findings indicate that enhanced monomer heterogeneity, and higher average CCS values, correlates with increased aggregation propensity.

**Figure 5.**
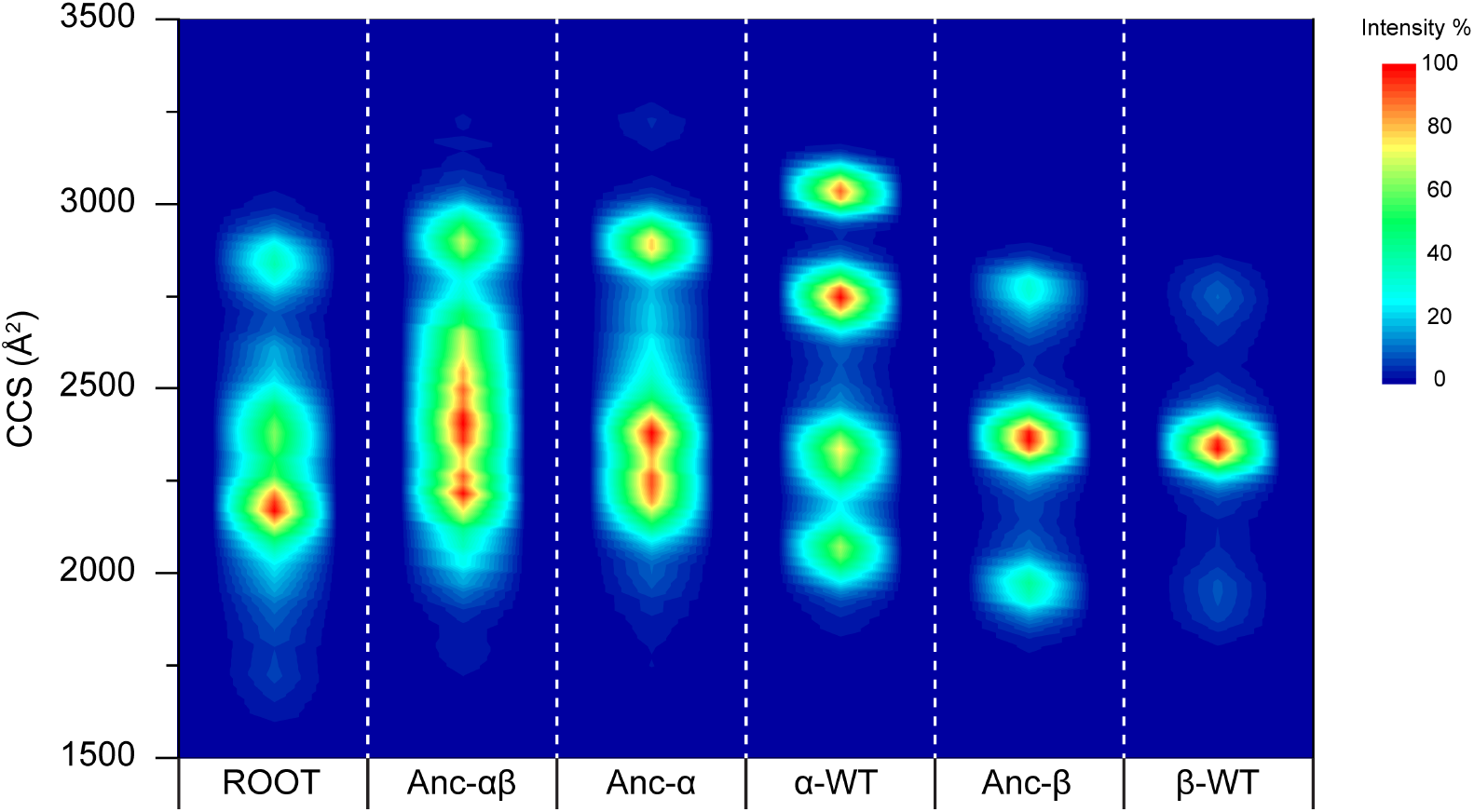
Evolution of synuclein monomer conformational diversity and collision cross section by ESI-IMS. Collision cross section (CCS) profiles for the compact +8 ion for each ancestral protein and for α-WT and β-WT measured by IMS. Data was collected over five minutes and averaged. All experiments were conducted at 20 µM protein concentration in 20 mM ammonium acetate (pH 7.4). CCS values were calculated using a calibration curve, detailed in methods.

### Ion mobility collision induced unfolding (CIU) and NMR paramagnetic relaxation enhancement (PRE) experiments reveal enhanced stability and localized intramolecular contacts in monomeric ROOT synuclein

To understand the conformational and dynamic differences between ROOT and α-WT, we used ion mobility collision induced unfolding (CIU) and performed paramagnetic relaxation enhancement (PRE) NMR experiments, to provide complementary views of protein stability and transient intramolecular monomer interactions. We assessed conformational stability using CIU, which monitors how protein ions respond to increasing collisional energy. In CIU, ions are accelerated into the IMS cell using incrementally higher collision voltages, where collisions with inert buffer gas induce unfolding of compact species^27^. Conformations observed at lower voltages reflect intrinsically more stable structures, whereas species appearing only at higher energies indicate less stable states. Our CIU analysis shows that compact ROOT is more stable than compact α-WT (Figure 6). Compact α-WT species begin to unfold at 10–15 V, while ROOT compact species remain largely intact until ≥25 V. Beyond these thresholds, both proteins rapidly lose all but the most extended conformers, indicating that the ROOT compact monomer is highly stable compared with α-WT.

**Figure 6.**
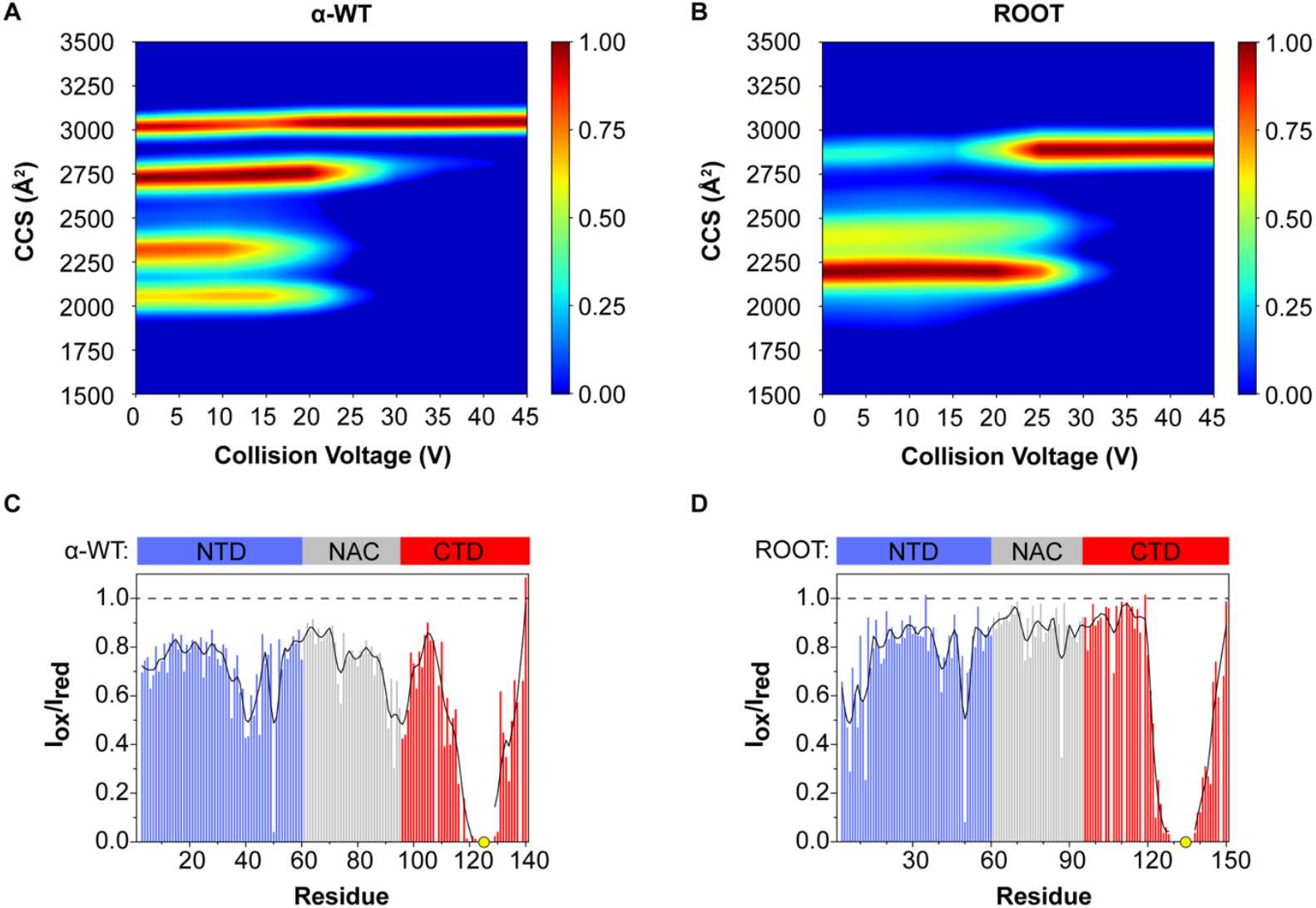
ROOT ancestor has stable compact species with enhanced N-C-terminal interactions. CIU heat maps showing the collision cross section for the conformational distribution of the +8 fingerprint species for α-WT (A) and ROOT (B) monomer at trap collision energy voltages ranging from 0-45V. All samples for CIU were at a concentration of 20 μM in 20 mM ammonium acetate (pH 7.4). (C-D) Intramolecular PRE intensity ratios (oxidized/reduced) for α-WT and ROOT synuclein. Yellow circles indicate location of MTSL spin label at position A124C for α-WT and A133C for ROOT. Dashed line at the PRE intensity ratio of 1 is included for reference. All PRE samples were composed of 200 μM protein in NMR buffer (20 mM phosphate, 100 mM NaCl, pH 6.8) with 10% D_2_O. Blue, grey, and red bars indicate the NTD, NAC, and CTD respectively.

To probe the molecular basis of this stability, we used PRE NMR experiments to examine transient long-range intramolecular interactions. PRE experiments introduce paramagnetic labels at specific sites, enabling detection of regions that transiently interact and providing insight into conformational ensembles and dynamic contacts often invisible to conventional structural techniques. The 2D ^1^H-^15^N HSQC spectrum for ROOT synuclein (Figure S9) shows narrow chemical shift dispersion typical of an IDP. NMR assignments were obtained for 143 non-proline residues, except for residues M1 and D2. Site directed spin labeling for PRE experiments, using 1-Oxyl-2,2,5,5-tetramethyl-3-pyrroline-3-methylmethanethiosulfonate (MTSL) as the paramagnetic spin label, were introduced at four positions across the NTD, NAC, and CTD. To minimize positional bias due to the differing lengths of the proteins, we selected four positions for spin-label incorporation corresponding to equivalent positions in both proteins. The selected positions for α-WT (A19, A90, A124, and A140) and ROOT (A19, A90, A133, A150) are representative of each domain and have been used in previous PRE experiments.

A general trend that is observed is that labeled MTSL sites (Figure 6, S9) in α-WT display a broader range of transient contacts across the monomer (I_ox_/I_red_ ∼0.8), whereas ROOT maintains higher values (I_ox_/I_red_ ∼0.9–1.0), highlighting its reduced propensity for heterogeneous long-range interactions. More specifically, in α-WT, the A124C label shows enhanced interactions with the N-terminal domain (NTD, residues ∼35–45) and NAC region (residues ∼90–100), contacts absent at the equivalent ROOT site (A133C). Conversely, ROOT A133C and A150C (Figure S9) labels show increased contacts with the extreme NTD (residues ∼1–12 and ∼1–5, respectively), indicating more localized intramolecular interactions. Overall, the PRE data demonstrate that ROOT samples fewer long-range transient interactions, but adopts a more compact conformation with enhanced local NTD–CTD contacts. This is consistent with CIU results, where ROOT exhibits greater resistance to unfolding, suggesting that its limited transient contacts contribute to a more stable, compact monomer. Together, CIU and PRE provide a complementary picture in which ROOT is both more conformationally stable and dynamically constrained relative to α-WT.

## DISCUSSION

Protein aggregation is a hallmark of neurodegenerative diseases, yet the molecular basis by which sequence dictates αSyn fibril formation remains incompletely understood. ASR offers a novel approach to experimentally probe the evolutionary origins of aggregation, providing access to ancestral protein states that cannot be observed in modern sequences. By resurrecting and characterizing key ancestral synucleins, including ROOT, the last common ancestor of all synucleins, and intermediate nodes along the αSyn and βSyn lineages, we gain a unique perspective on how sequence and structural features of the monomer and the fibril evolved to promote αSyn aggregation.

Our evolutionary analysis indicates that aggregation is a derived trait. ROOT synuclein represents an aggregation-resistant ancestral state that is intrinsically disordered yet compact and conformationally homogeneous relative to extant α-WT. Although ROOT contains a conserved NTD, a full-length NAC, and a negatively charged CTD, features generally sufficient for aggregation, it resists fibril formation across varied pH and salt conditions. Sequence analyses suggest that additional charged and aromatic residues strengthen NTD–CTD intramolecular interactions, promoting compaction and stability. This model is supported by PRE data, which show enhanced NTD–CTD interactions compared to α-WT, and by CIU experiments confirming greater stability of compact ROOT. Together, these findings indicate that ROOT embodies a stable, aggregation-resistant ancestral conformation that preceded the evolutionary acquisition of fibril-forming capacity.

Our phylogenetic and biophysical analyses reveal that the transition from ROOT to Anc-αβ marks the evolutionary origin of synuclein aggregation. This transition reflects a shift in sequence composition and charge distribution that collectively remodel the conformational landscape of the monomer. Anc-αβ exhibits a broader distribution of conformers relative to ROOT, as shown by IMS, with populations spanning multiple compact states. This expansion of the conformational ensemble coincides with the first appearance of aggregation, suggesting that monomer conformational heterogeneity may underly the emergence of fibril-forming capacity by providing access to aggregation competent monomer conformations. Previous studies have suggested that extended, diverse conformers may facilitate NAC exposure and fibril nucleation^28^, whereas highly compact and homogeneous monomers may not be aggregation competent^29^. Subsequent evolution in the αSyn lineage shows further expansion of monomer conformational diversity, while the βSyn lineage retains a compact ensemble and remains aggregation-incompetent under physiological conditions. The patterns observed in the ancestral lineage suggest that fibril formation depends not merely on monomer size or compaction, but on the interplay between conformational stability and heterogeneity. Collectively, our findings indicate that the evolutionary trajectory from ROOT to extant synucleins reflects a progressive tuning of monomer conformational ensembles, which may serve as a central determinant of synuclein aggregation propensity.

The emergence of aggregation along the αSyn lineage was further shaped by evolutionary tuning of the N-terminus, which rapidly evolved aggregation enhancing residues. Key substitutions in the P1 region (residues 36-42) accelerated fibril formation and importantly are conserved along the αSyn lineage, aligning with previous work identifying this region as a master regulator of aggregation^30^. Residues 38 and 42 in the P1 region of the NTD have been shown to be critical: S42A abolishes fibril formation, whereas L38A has little effect on kinetics; the L38M mutation (as in γ-WT) suppresses α-WT aggregation, while L38I accelerates it^31^. Furthermore, mutations such as M38I in γ-WT have been linked to amyloid deposits in patients suffering from amyotrophic lateral sclerosis^32^. Notably, our chimera experiments with ROOT and Anc-αβ demonstrate that the M38L and T42S substitutions in the Anc-αβ N-terminus accelerate aggregation, emphasizing the evolutionary tuning of the NTD as an early step in modulating fibril-forming propensity.

The transition from ROOT to α-WT is marked by the stepwise acquisition of residues critical for stabilizing the fibril core, initially enriching residues that form and stabilize the bent β-arch^33^, followed by substitutions that stabilize protofilament interactions^33^ (Figure 7). In α-WT fibrils, the structured core typically spans residues 36–99, encompassing portions of the NTD and CTD and the entire NAC^34^. Three regions are central to aggregation propensity: the preNAC (residues 47–56), forming the protofilament interface in rod polymorphs; the NACore (residues 68–78), mediating protofilament contacts in twisted polymorphs; and the β-arch^33,34^. The sequence sensitivity of these motifs is underscored by familial mutations that have been shown to remodel protofilament architecture^35-38^, highlighting their central role in fibril assembly. The evolution from ROOT to Anc-αβ coincides with key mutations in the NAC region such as K79Q with E86G and V65Q with E68G introduce hydrogen-bonding pairs analogous to those in α-WT, while S72T and N75T add branched hydrophobic side chains that reinforce core packing. These changes enable Anc-αβ fibril formation, consistent with chimera experiments showing that constructs with the Anc-αβ NAC aggregate, whereas those with the ROOT NAC constructs do not (Figure 4). Even subtle differences proved decisive, as shown with chimera experiments: αβαβR formed fibrils, whereas αβRR did not, underscoring the fine sequence sensitivity of the NAC region. Anc-αβ to Anc-α substitutions further stabilize the fibril core and optimize protofilament interactions, including S54T, A63V, while the Anc-α to α-WT transition adds S64T and Q65N, the latter forming a stabilizing hydrogen bond with G68 that strengthens the fibril fold. This stepwise trajectory illustrates how aggregation propensity arose incrementally, with the first evolutionary stage allowing for fibril formation and each subsequent evolutionary stage reinforcing the fibril architecture. This framework provides an evolutionary perspective on synuclein aggregation not accessible through traditional structural or biochemical approaches.

**Figure 7.**
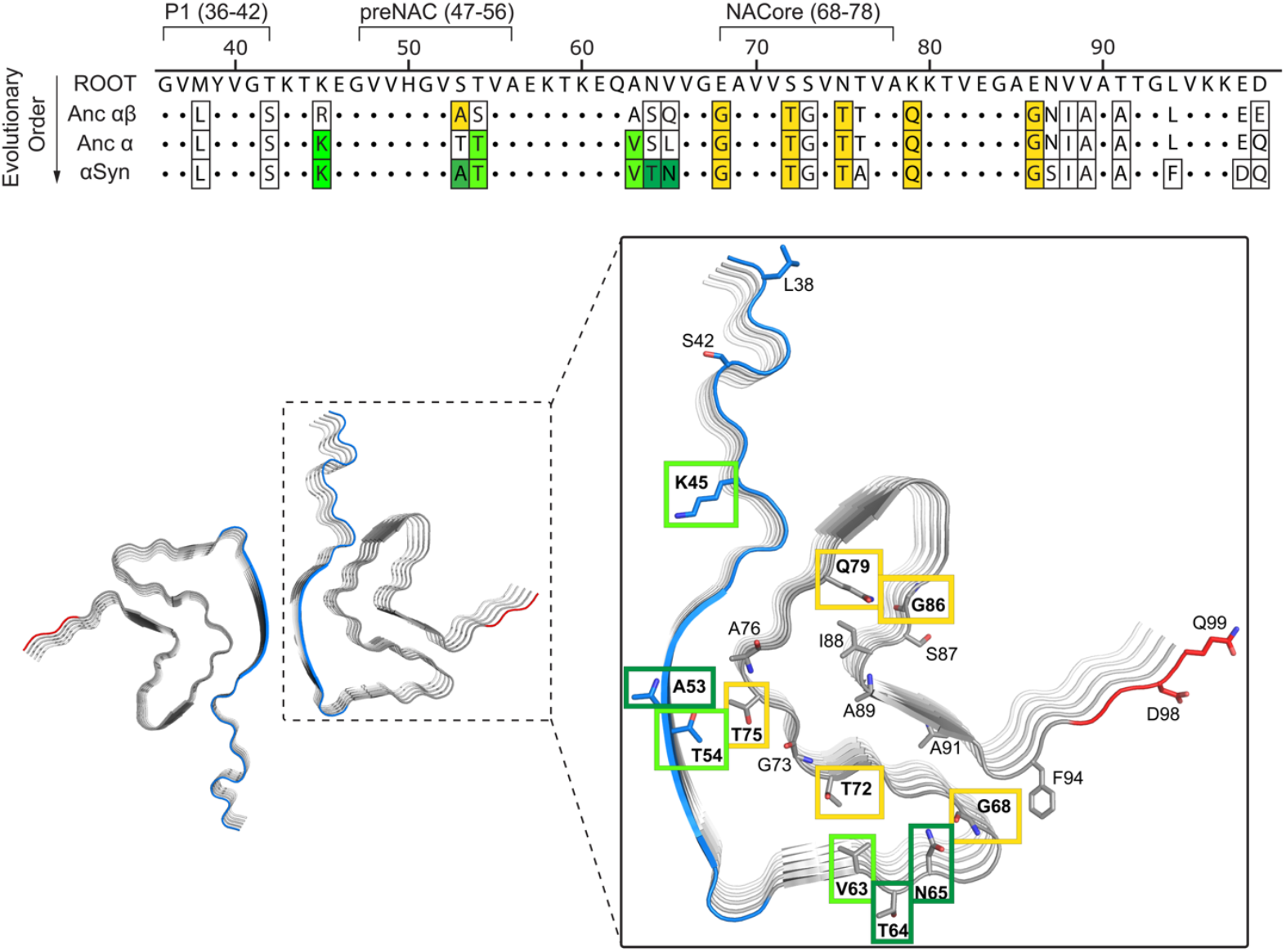
From ROOT to α-WT: Stepwise evolutionary acquisition of fibril-stabilizing residues in synuclein. Top: MSA of the fibril core (residues 36-99) for ROOT, Anc-αβ, Anc-α, and extant α-WT highlighting sequence changes that enabled fibril formation. Conserved residues are indicated by dots. Mutations relative to ROOT sequence are boxed, with colored boxes highlighting substitutions critical for forming and stabilizing the PBD: 6A6B fibril core. Yellow indicates mutations arising in Anc-αβ, bright green in Anc-α, and dark green in α-WT. Bottom: Schematic representation of evolution of fibril stabilizing residues mapped onto fibril structure of the α-WT “rod” polymorph (PBD: 6A6B). ROOT lacks key stabilizing features and remains aggregation-resistant and progressive substitutions define a stepwise evolutionary trajectory in which β-arch stabilization emerges first (yellow boxes), followed by optimization of protofilament-protofilament interactions (bright green), culminating in further fibril stabilization in α-WT (dark green). The zoomed-in view shows a single fibril protofilament with side chains displayed for all residues that differ from ROOT. Boxed residues correspond to those highlighted in the alignment above. The domains are color coded: blue: NTD, dark grey: NAC, Red: CTD.

## CONCLUSION

Our findings identify the ROOT → Anc-αβ transition as a singular evolutionary turning point at which two enabling features appeared simultaneously to give rise to synuclein aggregation: increased monomer conformational heterogeneity and fibril-stabilizing substitutions within the NAC that stabilize the β-arch core (Figure 8). The co-emergence of these traits provides both conformational access to aggregation-competent monomers and the structural determinants required for stable fibril architecture. Strategically positioned sequence changes transformed ROOT from an aggregation-resistant ancestor into an aggregation-prone protein at Anc-αβ, setting the stage for the divergent fates of the α- and β-synuclein lineages. Subsequent diversification within the αSyn and βSyn lineages fine-tuned these features, enhancing aggregation in αSyn through expanded conformational diversity and optimized fibril contacts or suppressing aggregation in βSyn by maintaining compact monomers and deleting a portion of the NAC sequence. This integrated view demonstrates that neither monomer conformational dynamics nor fibril-stabilizing motifs alone are sufficient for fibril formation; rather, it is their combined emergence that defined the evolutionary origin of synuclein fibril formation (Figure 8). By uniting sequence evolution with structural dynamics, our work offers a molecular basis for understanding how synuclein aggregation arose and establishes a generalizable paradigm for probing the evolutionary roots of protein aggregation in disease. Our study offers conceptual and mechanistic insights that may guide strategies to stabilize non-aggregating conformations or disrupt key aggregation motifs, ultimately advancing efforts to intervene in neurodegenerative disease.

**Figure 8.**
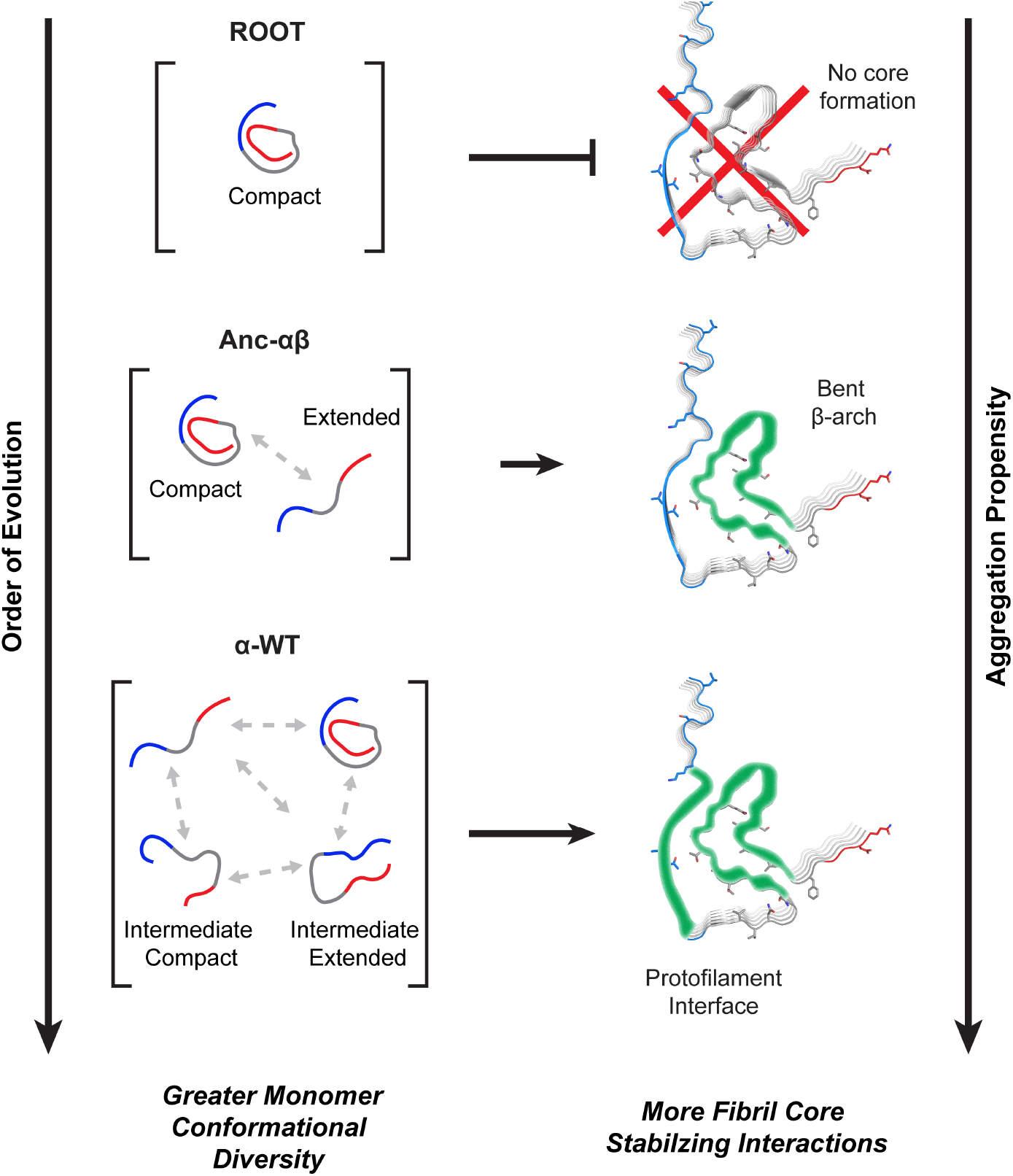
ASR reveals the stepwise evolution of aggregation drivers: greater monomer conformational diversity and increased fibril stabilizing interactions in NAC. At the monomer level (left), ROOT synuclein adopts a compact and conformationally homogeneous ensemble, Anc-αβ samples a broader range of conformations, and α-WT exhibits the greatest conformational diversity. At the fibril level, substitutions in Anc-αβ introduce intramolecular contacts that stabilize the bent β-arch core, while additional substitutions in α-WT further stabilize the core and refine the protofilament–protofilament interface. Together, the evolutionary emergence of more complex monomer ensembles and increasingly stabilizing interactions within the fibril promote aggregation-prone α-synuclein.

## MATERIALS AND METHODS

### Synuclein dataset compilation and refinement by simple sequence removal criteria

Sequences of αSyn, βSyn, and γSyn were compiled from the NCBI Protein database to form a preliminary dataset^39^. The NCBI Protein database was parsed with an in-house python script using the Biopython implementation of the Entrez Global Query Cross-Database Search System^40^ and synuclein sequences were identified using the gene search parameter with keywords “SNCA”, “SNCB”, or “SNCG”. The search yielded a starting dataset of 2583 synuclein sequences when queried on February 18, 2022. Initial dataset processing was performed with an in-house python script. Dataset entries with replicate sequences were removed to ensure that each entry was unique. Likewise, entries annotated as “incomplete” or lacking an annotation indicating a synuclein sequence (i.e. lacking “synuclein” or a synuclein alias in its sequence name or description) were also removed. Finally, because downstream phylogenetic inference lacks built-in capacity for accurately differentiating insertions from deletions during ASR, sequence length was used as an approximation for identifying and removing sequences with lengthy insertions or deletions likely due to sequencing errors, aberrations, and alternative splicing. Within each subfamily, the length of the sequences was strongly centered on the mean sequence length, so all sequences from a particular subfamily that exceeded ±1 standard deviation of the subfamily mean were removed. The sequence length means were 137.5 for αSyn, 130.1 for βSyn, and 133.1 for γSyn. Preliminary refinement resulted in a dataset of 783 unique synuclein sequences.

### Refinement of dataset by unique k-mer removal criteria and single isoform selection

Although sequence length has previously been used as a quick method for dataset processing in preparation for phylogeny inference^22^, accurate ASR requires more precise removal of lengthy insertions and deletions and selection of at most, a single isoform per species per ortholog. As such, we developed a dataset refinement protocol based on the removal of sequences containing unique k-mers.

Unique k-mer dataset refinement relies on two criteria for determining sequences to be discarded. The first criterion, gap occupancy (*g*), is defined as the percentage of sequences containing a gap in a given position of a multiple sequence alignment (MSA) of the dataset. The second criterion, unique k-mer length (*k*), is defined as the length of a continuous and completely unique subsequence within a sequence of the MSA of the dataset. Because insertions and deletions that would inform the ASR of the most ancestral nodes of a phylogeny must have been propagated through several sequences in a sufficiently large dataset, discarding only sequences with unique insertions and modest gap occupancy should considerably optimize the dataset for ASR. An in-house python script was developed to implement unique k-mer dataset refinement, utilizing an initial unique k-mer length *k*=2 and no gap occupancy criterion to focus on discarding alternative isoforms. Refinement was performed repeatedly on the Rutgers Amarel research computing cluster (Office of Advanced Research Computing at Rutgers, The State University of New Jersey), and after each successive round of refinement, the MSA was recalculated using Clustal Omega^41^ until no further refinement could be achieved, producing a dataset of 692 synuclein sequences. Subsequently, isoforms were discarded until each species had no more than a single isoform for each synuclein. Following isoform selection, the unique k-mer refinement was repeated with a gap occupancy *g*=0.33 and unique subsequence length *k*=2, which resulted in the final dataset of 603 synuclein sequences.

### Phylogeny inference, ancestral sequence reconstruction and sequence analysis

A synuclein phylogeny was inferred from the final dataset using Randomized Accelerated Maximum Likelihood (RAxML)^20^ rapid bootstrap analysis to search for the best-scoring maximum likelihood (ML) tree. The MSA of the final dataset was used as the input alignment for the phylogeny inference. The ML tree was subsequently rooted using the RAxML algorithm for balanced subtree branch length tree rooting. Finally, the sequences for each ancestral node in the ML tree were determined using the RAxML algorithm for computing marginal ancestral states. All computations using RAxML algorithms were performed with a disorder-specific amino acid substitution scoring matrix^42^ and the Gamma model of mutation rate heterogeneity^43^. An MSA of key extant and ancestral sequences was performed using Clustal Omega. A consensus sequence was calculated from the MSA with a minimum consensus threshold of greater than 50% and the alignment was visualized using SnapGene Viewer.

### Plasmid production, protein expression, and protein purification

Plasmids encoding the relevant sequences were designed and inserted into the pT7-7 αSyn WT plasmid backbone (Addgene plasmid #36046) between the NdeI (5’) and HindIII (3’) sites and were ordered from Azenta Life Sciences. Plasmids were transformed into *E. coli* BL21(DE3) chemically competent cells (Invitrogen Inc.) and plated with appropriate antibiotics. Protein expression and purification was performed as previously described^44^, with Anc-α and α-WT elution at 50% elution buffer, while all other synucleins were eluted at 60% elution buffer. Isotope labeled protein (^15^N;^13^C) was grown at the same conditions in M9 minimal media supplemented with ^15^NH_4_Cl and/or ^13^C-glucose.

Cysteine mutants were created at selected residues (A19C, A90C, A133C, and A150C in ROOT; A19C, A90C, A124C, and A140C in α-WT) by designing primers using the NEBaseChanger tool (NEB). Site directed mutagenesis was performed according to manufacturer’s instructions using the Q5 Site Directed Mutagenesis Kit (NEB). All plasmids were confirmed by Sanger sequencing (Genewiz).

All purified proteins were analyzed by SDS-PAGE or native electrospray ionization mass spectrometry (ESI-MS) using a SYNAPT G2-SIqTOF (Waters, MA, USA). All samples were flash-frozen in liquid nitrogen and stored at −80°C until further use.

### Solution state NMR

Purified ^15^N and/or ^13^C-labeled protein was concentrated and buffer exchanged into NMR buffer (20 mM phosphate, 100 mM NaCl, pH 6.8) and diluted to a final concentration of 350 μM before being supplemented with 10% D_2_O. ^1^H, ^15^N, and ^13^C assignments were indirectly referenced to 1 mM DSS^45^. All NMR backbone assignment experiments were performed at 298 K on a Bruker Avance III HD 800 MHz spectrometer. NMR backbone assignment experiments used include: 2D ^1^H-^15^N HSQC, 3D HNCO, 3D HN(CA)CO, 3D CBCA(CO)NH, 3D HNCACB, and 3D (H)N(COCA)NNH. All NMR data was processed using both Topspin 4.10 (Bruker) and NMR Pipe/NMRDraw^46^ and analyzed using Sparky^47^ and CARA^48^.

### Paramagnetic relaxation enhancement (PRE) NMR

Purified cysteine mutants were incubated with 1 mM DTT at 4°C for 3 hours. Excess DTT was removed using a desalting column (PD10, Cytiva) and the proteins were immediately combined with 5-fold molar excess of MTSL (1-Oxyl-2,2,5,5-tetramethyl-3-pyrroline-3-methylmethanethiosulfonate, Sigma) and incubated in the dark at 4°C for 14 h. Excess MTSL label was removed by concentrating and buffer exchanging the samples into NMR buffer using a 3kDa spin filter. Final solutions were diluted to 200 μM with NMR buffer and supplemented with 10% D2O. Spectra were acquired using an Avance III HD 800 MHz spectrometer equipped with a TCI cryo-probe at 15°C. The spectrum for the oxidized sample was obtained first, followed by reduction using ascorbic acid, then acquisition of the spectrum of the reduced sample. Samples were reduced by adding ascorbic acid to a final concentration of 2 mM, followed by incubation for 6 hours at room temperature. The acquisition parameters for the spectra were 64 scans and 2048 (F2) and 256 (F1) complex points. Spectral widths were 16.03 ppm (^1^H) and 35.00 ppm (^15^N), respectively. Peak heights were used to calculate the PRE intensity ratios (I_ox_/I_red_).

### Thioflavin T fluorescence assay

Protein aggregation was analyzed by use of a Thioflavin T (ThT) fluorescence. Protein samples were concentrated and buffer exchanged into PBS (pH 7.4) with a 3kDa centrifugal spin filter and aggregates were removed using a 50kDa spin filter. Final solutions were diluted to the appropriate concentrations (70 μM unless otherwise indicated) and were mixed with ThT. Samples were loaded into a 96-well plate, and each well was supplemented with a single 3 mm Teflon bead and ThT at a final concentration of 20 μM. Sealed plates were loaded into a POLAR Star Omega Plate Reader (BMG Labtech; NC, USA) and assays were conducted at 37°C with shaking at 600RPM with fluorescence intensity (480nm) measured every 33 minutes over 140 hours. Fluorescence traces were normalized to the final fluorescence intensity and replicates were averaged. Following completion of the assay, replicates were pooled and centrifuged at 15,000RPM for 2 hours to pellet fibrils. The supernatant was removed, and the concentration of residual soluble species was measured for each sample using a Nanodrop 2000. The extinction coefficient used was 5960 M^-1^cm^-1^ as calculated using the Expasy ProtParam tool. Averaged ThT curves were fitted to equation 1 below using OriginPro, where A_1_ and A_2_ are the base and maximum fluorescence respectively, x_0_ is the half time, and dx is the time constant. Lag times were calculated using equation 2 below.

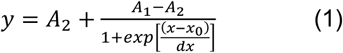

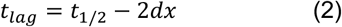

### Atomic force microscopy

Fibrils or insoluble aggregates obtained from ThT assay plates were dispersed in 200 μL of PBS and subsequently diluted 10-fold. The diluted solution was deposited onto a clean 1 cm x 1 cm mica surface and incubated at room temperature for 10 minutes. The surface was then washed four times with 1 mL of ultrapure water to remove excess salt and unbound proteins. The mica surface was allowed to air-dry for 1 hour prior to imaging. Freshly prepared mica samples were used for AFM imaging. AFM images were acquired in non-contact mode using a Cypher ES atomic force microscope (Asylum Research) equipped with PNPDB tips. All images were collected at room temperature with a resolution of 512 x 512 pixels. Acquired images were processed and analyzed using Gwyddion software.

### Native ESI-MS and assessment of charge state distribution

All native ESI-MS was performed with sample concentrations of 20 µM protein in 50 mM ammonium acetate (pH 7.4), collected over a 3-minute period using an in-house Synapt G2-SI qTOF at 3kV capillary voltage, 40V sample cone, and a source temperature of 100°C. Spectra were analyzed using the MassLynx software (Waters). The results from the native ESI-MS were used to create plots of charge vs. intensity for ions from +7 to +22, which were then fitted to Gaussian distributions to determine relative populations distributions of conformers in solution. Gaussian fitting was conducted using the Multiple Peak Fit tool in Origin Pro 8.0. Population percentages were determined by using the area for each fitted Gaussian.

### Ion mobility-mass spectrometry and collision induced unfolding experiments

Ion mobility-mass spectrometry (IMS) was conducted using a Synapt G2-SI qTOF, with samples at 20 μM concentration in 20 mM ammonium acetate (pH 7.4). IMS conditions include: 1.4kV capillary voltage, 40V sample cone, source temperature of 30°C, 4V trap CE, 500m/s IMS wave velocity, and 35V IMS wave height. IMS calibration was conducted using myoglobin, cytochrome C, and ubiquitin (Sigma Aldrich) using the accepted CCS values from the Bush Database^49^. Each standard was prepared at 10 µM in 50% acetonitrile, 50% water, 0.1% formic acid. All IMS runs were collected over 5 minutes and averaged. Drift time plots were extracted by using Driftscope software (Waters) and used with the calibration curve to determine the collision cross section (CCS). Collision-induced unfolding (CIU) experiments were conducted at 20 μM concentration in 20 mM ammonium acetate (pH 7.4). CIU conditions were the same as those used for regular IMS, but with a varying trap collision energy (Trap CE) ranging from 0-45V, which was incremented at 5V intervals per run. Each run was collected over 5 minutes and averaged.

### Statistical analysis

Data was analyzed via two-sample t-tests, assuming either equal variance or unequal variance where indicated. For samples with unequal variances, Welch’s correction was applied. Tests were conducted using OriginPro and results were considered significant for p<0.05 or better.

## Author Contributions

J.B designed the research; J.W and A.S. and E.L. performed computational ancestral sequence reconstruction and constructed phylogenetic tree; A.S and S.L. generated ancestral sequence plasmids, purified proteins, and performed ThT assays; S.L generated chimera plasmids and performed NMR PRE experiments; A.N. performed the solution state NMR assignments; J.E. performed sequence analysis of ancestral proteins, performed mass spectrometry experiments and analysis; A.S, J.E and J.B analyzed the data; A.S, J.E and J.B. wrote the manuscript.

## Acknowledgements

This work was supported by an NIH grant GM136431 to Jean Baum

