## Supplementary Figures for "Is synuclein aggregation a derived or ancestral trait? Ancestral sequence reconstruction uncovers stepwise evolution of synuclein aggregation"

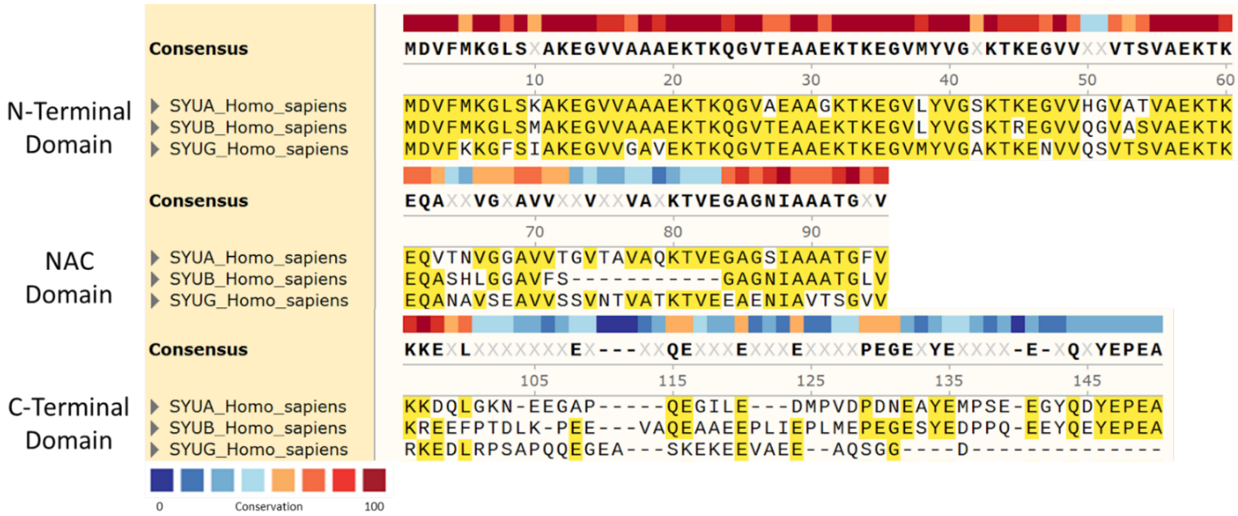


**Figure S1. Multiple sequence alignment of the full 603 sequence dataset for sequences of αSyn, βSyn, and γSyn.** Multiple sequence alignment (Clustal Omega) and corresponding consensus sequence derived from the full 603 sequence dataset. Only the extant human αSyn, βSyn, and γSyn sequences are shown in the alignment for brevity. Yellow boxes indicate residues that are the same as the indicated consensus sequence. Colored blocks above the consensus sequence represent conservation ranging from 0% (dark blue) to 100% (dark red). Dashes indicate gaps from MSA.

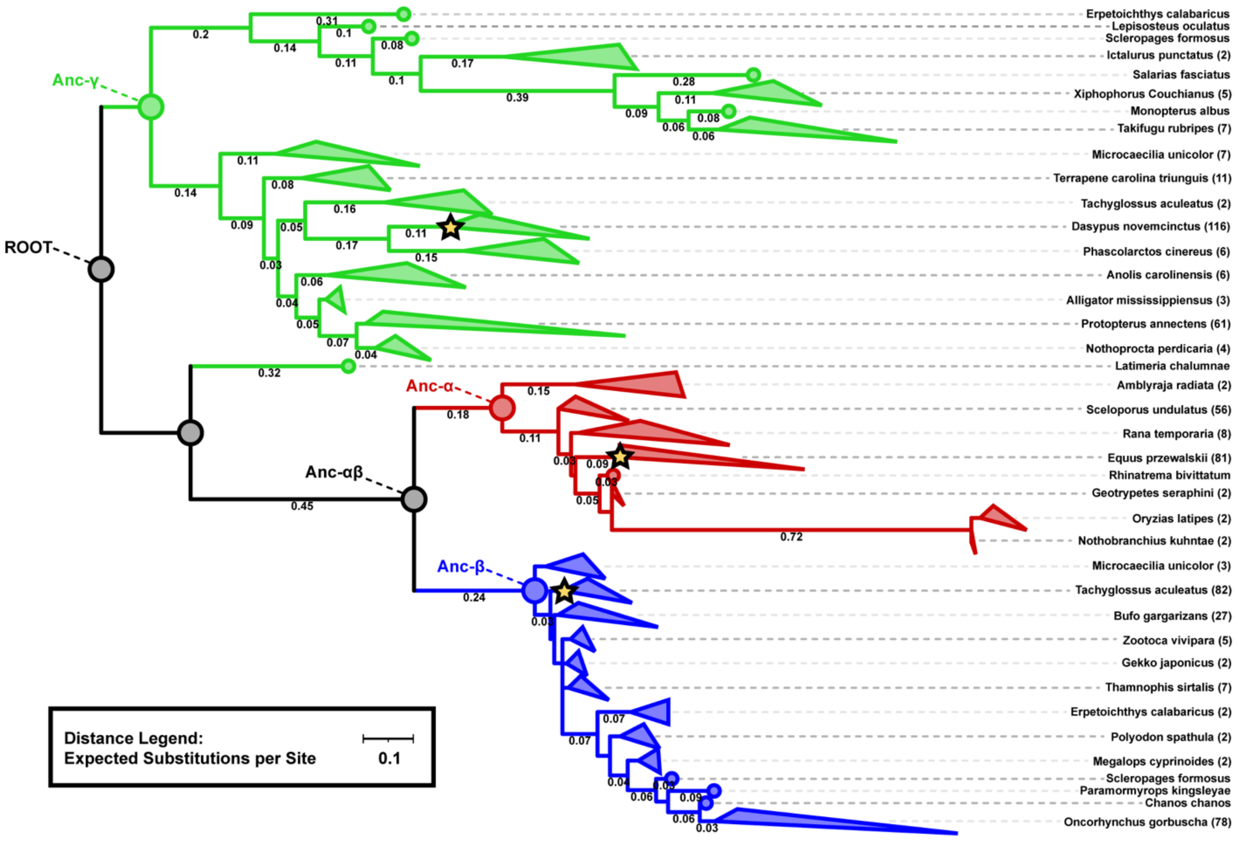


**Figure S2. Reconstructed synuclein phylogenetic tree.** A phylogenetic tree of 603 synuclein proteins created through an ancestral sequence reconstruction using Randomized Accelerated Maximum Likelihood (RAxML) software. The tree is colored according to synuclein clades, with α-synuclein clade represented in red, β-synuclein clade in blue, and γ-synuclein clade in green. The branch lengths correspond to genetic distance, with units of expected substitutions per site, or substitutions per protein residue. Branches including extant human synuclein sequences are indicated by yellow stars, and key ancestral nodes are indicated by large circles. Small circles represent individual extant sequences. Branches were collapsed into scalene triangles, where an average branch length less than 0.3 substitutions per site occurs. Any branch lengths less than 0.03 are not labelled. The size and shape of the scalene triangles show both the most closely related and most distantly related sequences within a given branch. The synuclein protein species names are aligned to the right, with the name of the most genetically distant species from the branch node and the total number of synucleins of the collapsed branch in parentheses.

**NTD**

| | ROOT | Anc-αβ | Anc-α | α-WT | Anc-β | β-WT |
|---|---|---|---|---|---|---|
| ROOT | 100 | 87 | 88 | 88 | 85 | 83 |
| Anc-αβ | | 100 | 93 | 95 | 98 | 95 |
| Anc-α | | | 100 | 98 | 92 | 88 |
| α-WT | | | | 100 | 93 | 90 |
| Anc-β | | | | | 100 | 97 |
| β-WT | | | | | | 100 |

% Identity: 80 | 90 | 100

**NAC**

| | ROOT | Anc-αβ | Anc-α | α-WT | Anc-β | β-WT |
|---|---|---|---|---|---|---|
| ROOT | 100 | 69 | 66 | 57 | 43 | 43 |
| Anc-αβ | | 100 | 94 | 83 | 60 | 57 |
| Anc-α | | | 100 | 86 | 54 | 54 |
| α-WT | | | | 100 | 46 | 46 |
| Anc-β | | | | | 100 | 96 |
| β-WT | | | | | | 100 |

% Identity: 40 | 50 | 60 | 70 | 80 | 90 | 100

**CTD**

| | ROOT | Anc-αβ | Anc-α | α-WT | Anc-β | β-WT |
|---|---|---|---|---|---|---|
| ROOT | 100 | 54 | 45 | 40 | 46 | 43 |
| Anc-αβ | | 100 | 69 | 45 | 73 | 64 |
| Anc-α | | | 100 | 67 | 45 | 43 |
| α-WT | | | | 100 | 38 | 39 |
| Anc-β | | | | | 100 | 92 |
| β-WT | | | | | | 100 |

% Identity: 30 | 40 | 50 | 60 | 70 | 80 | 90 | 100

| | NTD | | | NAC | | | CTD | | |
|---|---|---|---|---|---|---|---|---|---|
| | ⊕ | ⊖ | Net | ⊕ | ⊖ | Net | ⊕ | ⊖ | Net |
| ROOT | 12 | 8 | **+4** | 2 | 4 | **-2** | 3 | 19 | **-16** |
| Anc-αβ | 11 | 8 | **+3** | 1 | 2 | **-1** | 3 | 23 | **-20** |
| Anc-α | 11 | 7 | **+4** | 1 | 2 | **-1** | 3 | 18 | **-15** |
| α-WT | 11 | 7 | **+4** | 1 | 2 | **-1** | 3 | 15 | **-12** |
| Anc-β | 11 | 8 | **+3** | 0 | 1 | **-1** | 3 | 19 | **-16** |
| β-WT | 10 | 8 | **+2** | 0 | 1 | **-1** | 3 | 19 | **-16** |

| | NTD | | NAC | | CTD | |
|---|---|---|---|---|---|---|
| | $H_2O$ (crossed) | ↕ | $H_2O$ (crossed) | ↕ | $H_2O$ (crossed) | ↕ |
| ROOT | 22 | 11 | 16 | 10 | 19 | 9 |
| Anc-αβ | 24 | 10 | 16 | 10 | 17 | 9 |
| Anc-α | 23 | 11 | 17 | 9 | 17 | 13 |
| α-WT | 24 | 10 | 17 | 9 | 14 | 9 |
| Anc-β | 24 | 10 | 12 | 6 | 19 | 9 |
| β-WT | 24 | 11 | 12 | 6 | 19 | 8 |

Residue Key: ⊕ = Positive, ⊖ = Negative, $H_2O$ (crossed) = Hydrophobic, ↕ = Polar

**Figure S3. Pairwise sequence alignments and residue breakdown.** (Left) Pairwise sequence alignment diagrams for the N-terminal domain (top), NAC domain (middle), and C-terminal domain (bottom). Numbers indicate percentage identity for indicated pairs. (Right, top) Chart showing the charged residue composition for the ancestral and extant sequences, broken down by domain. (Right, bottom) Chart showing the residue type composition for the ancestral and extant protein sequences, broken down by domain. Legends are indicated at the bottom of the table. Hydrophobic residues correspond to: V, I, L, F, A, M, P. Polar residues: Y, S, T, Q, N, H.

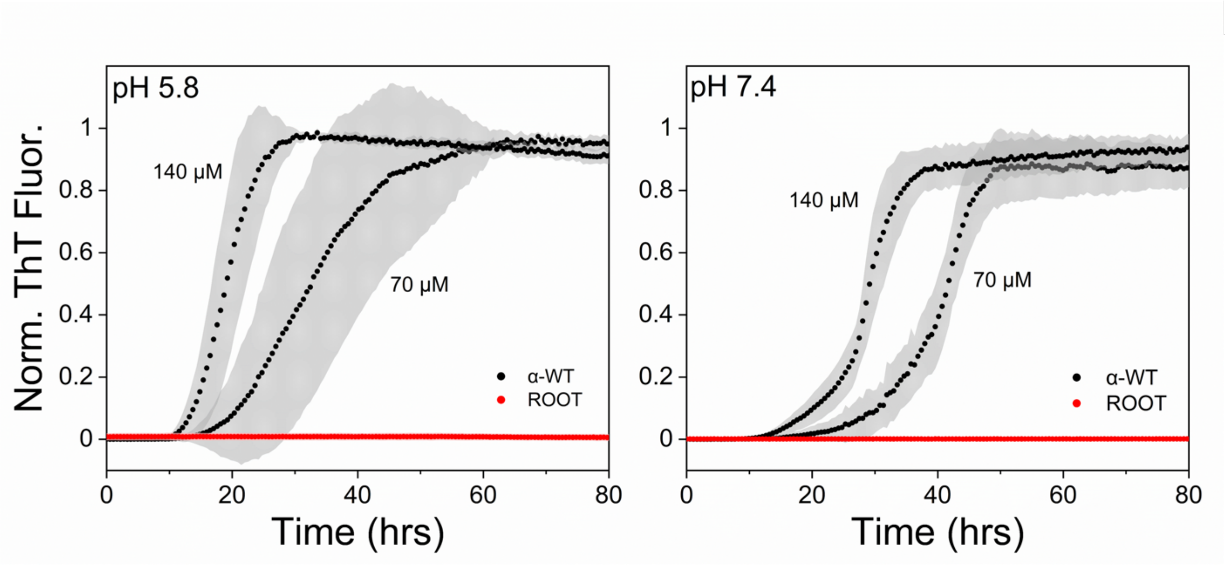


| pH | 5.8 | | 7.4 | |
|---|---|---|---|---|
| Concentration (μM) | 70 | 140 | 70 | 140 |
| ROOT Lag time (hrs) | - | - | - | - |
| α-WT Lag time (hrs) | 25.86 ± 0.56 | 15.07 ± 0.77 | 31.25 ± 0.30 | 22.46 ± 0.30 |

**Figure S4. ThT assay of ancestral ROOT at pH 7.4 and pH 5.8 reveal high resistance to aggregation relative to α-WT.** Top: Normalized ThT fluorescence assays for ancestral ROOT (red) and α-WT (black) at pH 5.8 and pH 7.4, at 70 μM and 140 μM. Traces represent average normalized fluorescence and standard deviation. The graph reaches a plateau with no signal change after 80 hours. Bottom: Table with α-WT lag times at different conditions calculated as described in Methods. ROOT synuclein has no measurable lag time in ThT experiment.

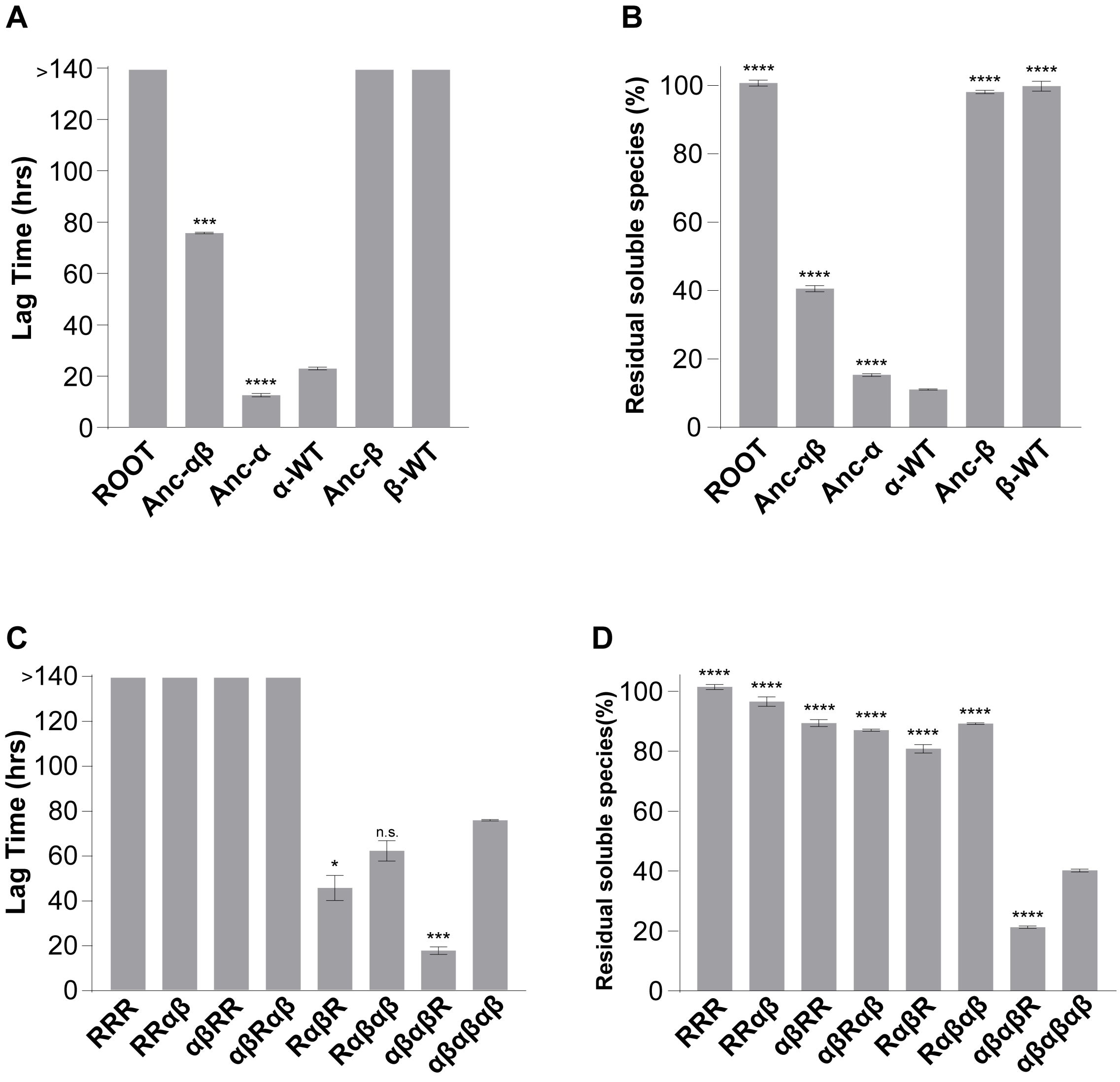


**Figure S5. Lag times and residual soluble species**. The average lag times (A) and residual soluble species (B) for the ThT assay shown in Figure 3. The average lag times (C) and residual soluble species (D) for the chimera ThT assay shown in Figure 4. Bars in (A,C) >140 hours had no measurable lag time. The average residual soluble species was calculated as a percentage of the initial protein concentration. Error bars represent standard deviation of at least 3 replicates. Significance was assessed by comparing each species to α-WT (A,B) or Anc-αβ (C,D) using an unpaired t-test. *$p<0.05$; **$p<0.005$; ***$p<0.0005$; ****$p<0.0001$.

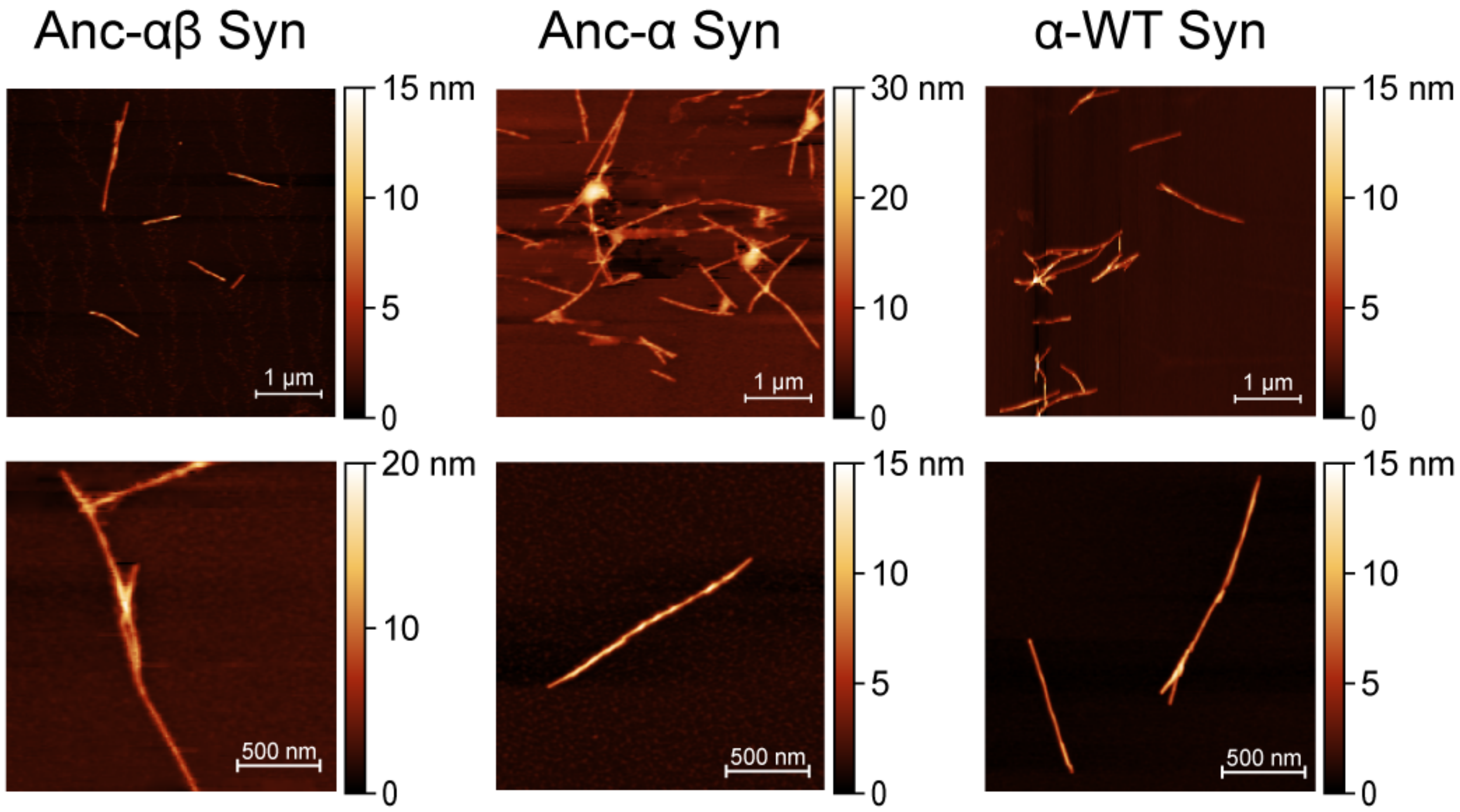


**Figure S6. AFM images of ancestral proteins and α-WT.**
AFM images of fibrils taken from endpoint ThT assays for Anc-αβ (left), Anc-α (middle), and α-WT (right). Top row images are a 1 μm while bottom rows are zoomed in at 500 nm.

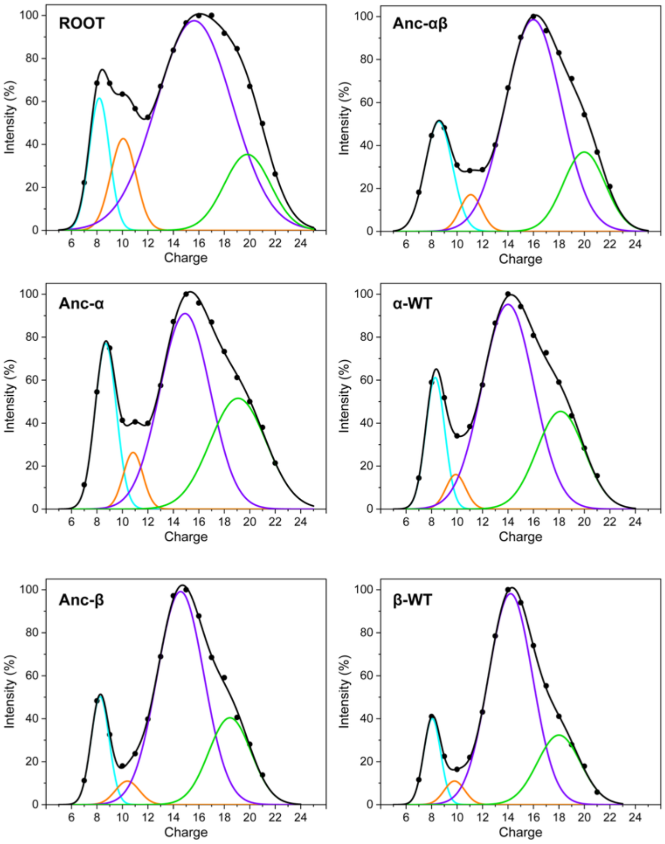


| ROOT | |
|---|---|
| *Main Charge State* | *Area % (SD)* |
| 8.2 | 11.0% (±0.6) |
| 10.1 | 9.4% (±0.8) |
| 15.6 | 65.2% (±1.5) |
| 19.8 | 14.4% (±0.9) |

| Anc-αβ | |
|---|---|
| *Main Charge State* | *Area % (SD)* |
| 8.6 | 15.8% (±0.4) |
| 11.1 | 4.2% (±0.3) |
| 16.0 | 62.5% (±0.6) |
| 20.0 | 17.5% (±0.4) |

| Anc-α | |
|---|---|
| *Main Charge State* | *Area % (SD)* |
| 8.7 | 17.0% (±0.4) |
| 10.8 | 5.4% (±0.3) |
| 14.9 | 47.6% (±0.6) |
| 19.1 | 30.0% (±0.5) |

| α-WT | |
|---|---|
| *Main Charge State* | *Area % (SD)* |
| 8.3 | 13.7% (±0.9) |
| 9.9 | 3.7% (±0.9) |
| 14.0 | 57.5% (±1.8) |
| 18.1 | 25.1% (±1.3) |

| Anc-β | |
|---|---|
| *Main Charge State* | *Area % (SD)* |
| 8.3 | 12.1% (±0.4) |
| 10.4 | 3.5% (±0.4) |
| 14.6 | 61.7% (±0.9) |
| 18.4 | 22.7% (±0.6) |

| β-WT | |
|---|---|
| *Main Charge State* | *Area % (SD)* |
| 8.0 | 10.2% (±0.3) |
| 9.8 | 3.4% (±0.3) |
| 14.2 | 65.4% (±0.7) |
| 18.0 | 21.0% (±0.5) |

**Figure S7. ROOT conformational distribution is unique and shows increased intermediate compaction relative to other ancestral node proteins.** Population distribution of charge states for each monomer are shown with fitted Gaussians for each distribution. The tables at the right detail the main charge states and the corresponding relative area % with standard error for each respective protein. Four main conformers are observed via native ESI-MS for each protein: compact (cyan), intermediate compact (orange), intermediate extended (purple), extended (green). All experiments were conducted at 20 μM protein concentration in 20 mM ammonium acetate (pH 7.4).

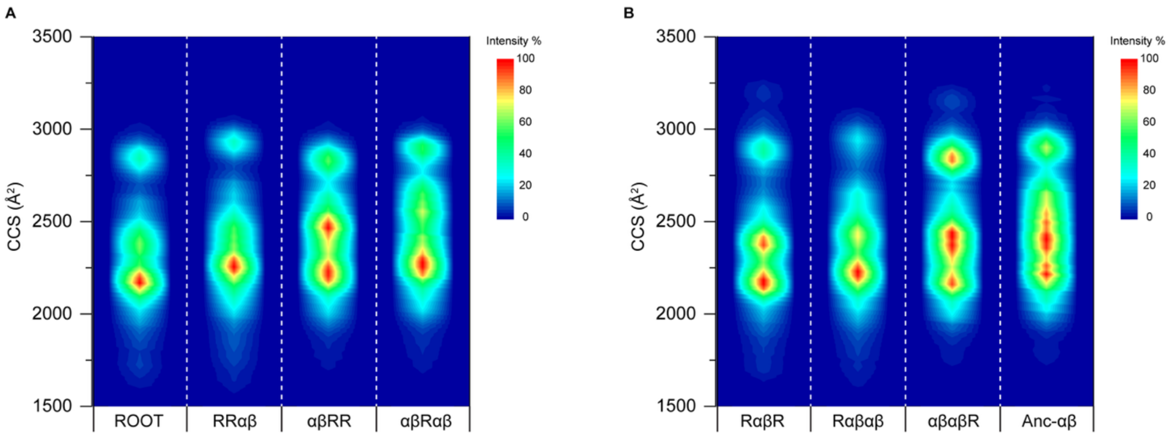


**Figure S8. Native IMS profiles for compact conformers of chimeric synucleins.** Collision cross section (CCS) heat maps for the +8 ion of each species, as measured by IMS. (A) CCS profiles for the +8 ion of ROOT and ROOT NAC chimeras. (B) CCS profiles for the +8 ion of Anc-αβ and Anc-αβ NAC chimeras. All experiments were conducted at 20 μM protein concentration in 20 mM ammonium acetate (pH 7.4).

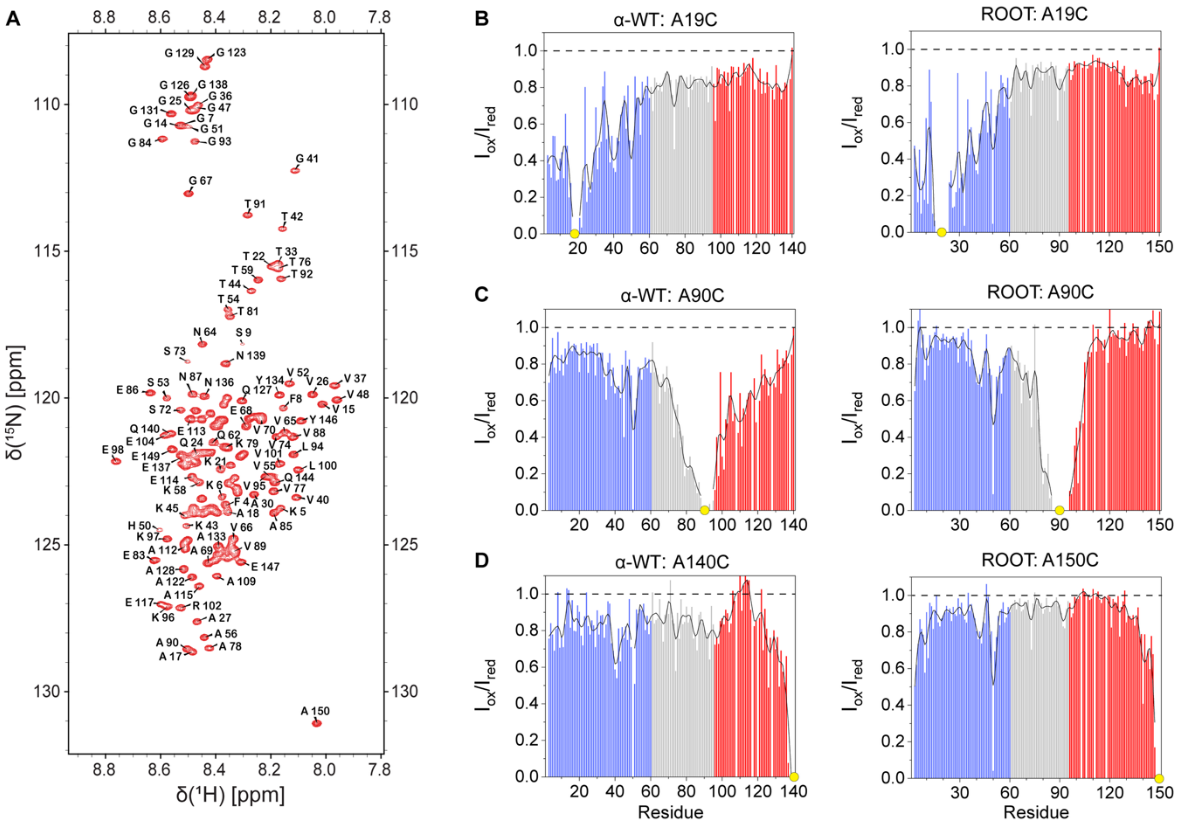


**Figure S9. NMR characterization of ROOT synuclein.** (A) 2D $^{1}H$-$^{15}N$ HSQC spectra of ROOT at pH 6.8 and 15°C, with residues labeled. (B-D) NMR PRE experiments for α-WT (left) and ROOT (right), with MTSL tags at A19C (top row), A90C (middle row), and A124C/A133C (bottom row). Dashed line at the PRE intensity ratio of 1 is included for reference. All samples were composed of 200 µM protein in NMR buffer (20 mM phosphate, 100 mM NaCl, pH 6.8) with 10% $D_2O$. Blue, grey, and red bars indicate the NTD, NAC, and CTD respectively.
